# A novel SARS-CoV-2-derived infectious vector system

**DOI:** 10.1101/2024.12.16.628561

**Authors:** Ghada Elfayres, Yong Xiao, Qinghua Pan, Chen Liang, Benoit Barbeau, Lionel Berthoux

## Abstract

Severe acute respiratory syndrome coronavirus 2 (SARS-CoV-2) is the causative agent of COVID-19. The development of antiviral drugs for COVID-19 has been hampered by the requirement of a biosafety level 3 (BSL3) laboratory for experiments related to SARS-CoV-2, and by the lack of easy and precise methods for quantification of infection. Here, we developed a SARS-CoV-2 viral vector composed of all four SARS-CoV-2 structural proteins constitutively expressed in lentivirally transduced cells, combined with an RNA replicon deleted for SARS-CoV-2 structural protein genes S, M and E, and expressing a luciferase-GFP fusion protein. We show that, after concentrating viral stocks by ultracentrifugation, the SARS-CoV-2 viral vector is able to infect two human cell lines expressing receptors ACE2 and TMPRSS2. Both luciferase activity and GFP fluorescence were detected, and transduction was remdesivir-sensitive. We also show that this vector is inhibited by three type I interferons (IFN-I) subtypes. Although improvements are needed to increase infectious titers, this vector system may prove useful for antiviral drug screening and SARS-CoV-2-related investigations.

## Introduction

The emergence of severe acute respiratory syndrome coronavirus 2 (SARS-CoV-2) in late 2019 marked the onset of a global health crisis unparalleled in recent history (Zhu et al., 2020). SARS-CoV-2 possesses a single-stranded positive-sense RNA genome of approximately 30 kilobases in length, encoding for a repertoire of proteins essential for viral replication, transcription, assembly and immune evasion (Kim et al., 2020). SARS-CoV-2 structural proteins include the spike (S), envelope (E), membrane (M) and nucleocapsid (N) proteins, which play pivotal roles in viral entry and assembly (Astuti, 2020). Notably, the S protein assumes a crucial role in the virus life cycle, enabling entry into human cells through interaction with the ACE2 receptor. Activation of the S protein is triggered by cellular proteases, including transmembrane protease serine 2 (TMPRSS2), further facilitating viral entry into host cells (Hoffmann et al., 2020). Advancement in vaccines and pharmacological agents against SARS-CoV-2 is still needed to address not only the COVID-19 pandemic but also future coronaviral outbreaks. Diverse vaccine formulations have emerged and received emergency authorization, leveraging platforms encompassing mRNA, viral vector, and protein subunit technologies (Polack et al., 2020;Sadoff et al., 2021;Vohra-Miller and Schwartz, 2022). However, the protection offered by these injectable vaccines has been consistently short-lived, and there is consensus that novel vaccines that can be delivered to the airways represent a more promising avenue (Dotiwala and Upadhyay, 2023;Ye et al., 2023). Antiviral drugs such as remdesivir and Paxlovid have been used to treat SARS-CoV-2. Remdesivir, an RNA polymerase inhibitor, was among the first antivirals authorized for emergency use (Beigel et al., 2020). However, its clinical effectiveness was controversial, with studies showing mild reductions in recovery time and little to no impact on overall mortality (Wang et al., 2020). Paxlovid, a combination of nirmatrelvir and ritonavir, has shown more promising results in early treatment of COVID-19, especially in high-risk patients, by reducing the severity of symptoms and the incidence of hospitalization (Niraj et al., 2022). However, no antiviral has been widely adopted due to limitations in their effectiveness, high costs, and logistical challenges of administration (Indari et al., 2021). Additionally, Paxlovid has raised concerns over post-treatment viral rebound, where patients experience a renewal of viral replication and symptoms after completing the treatment course, further limiting the utility of these antivirals as effective treatments (Parums, 2022).

SARS-CoV-2 antiviral research presents several challenges. Due to airborne transmissions and its potential for severe disease, many countries classified SARS-CoV-2 as a Biosafety Level 3 (BSL-3) pathogen. BSL-3 facilities are not universally accessible and greatly increase costs (Kaufer et al., 2020). In addition, the large size of coronaviral genomes (≈30 kb) complicates reverse genetics manipulations (Amarilla et al., 2021). Developing SARS-CoV-2-based vector systems with reduced biosafety risk has been an important goal (Ghosh et al., 2020). Furthermore, such vectors should ideally encode marker genes to allow for sensitive, easy and inexpensive quantification of infection yields (Jiang et al., 2008). Several SARS-CoV-2 model systems have been devised as substitutes for the wild-type virus and contain a varying number of SARS-CoV-2 genes (Chen and Zhang, 2021). Lentiviral vectors pseudotyped with the SARS-CoV-2 spike protein were created, but they can only be applied to the study of the cellular entry step, such as the evaluation of neutralizing antibodies (Condor Capcha et al., 2021). Other teams constructed SARS-CoV-2-based vector systems that rely on this its structural proteins and thus more truly reflect a coronaviral infection. All these systems are based on the complementation approach: they involve the introduction of a subgenomic RNA devoid of some of the structural proteins into mammalian cells, and the concomitant expression of the deleted structural proteins from other nucleic acids; vector particles that can perform a single cycle of infection are collected from the supernatants (Thiel et al., 2003). In what is perhaps the most minimalistic approach for such a model, Syed *et al*. created a system in which plasmids encoding the four SARS-CoV-2 structural proteins (S, E, M, N) are co-transfected, along with a messenger RNAs (mRNAs) expressing reporter proteins that also contains virion encapsidation signals (Syed et al., 2021). Although this system presents distinct advantages such as its versatility, it is limited by the inherently short duration of marker protein expression; in addition, since no viral RNA is present and no RNA replication takes place, it cannot be used to study post-entry replication steps. More recently, another team developed a similar system, again involving the co-transfection of the four SARS-CoV-2 structural proteins as well as a reporter protein-expressing mRNA coupled with the SARS-CoV-2 packaging signal sequence, but this RNA also included amplification elements derived from the Venezuelan equine encephalitis virus (VEEV), increasing duration of marker expression (Liu and Liu, 2023). Other teams have created SARS-CoV-2 vector systems that include the viral replication machinery, allowing for the study of post-entry viral replication steps. Ju et al. have reported a simple system whereby a SARS-CoV-2 replicon RNA in which the N protein is replaced by GFP, is able to propagate in cells stably expressing N; SARS-CoV-2 vectors produced by these cells can achieve single cycle infection of cells not expressing N (Ju et al., 2021). Ricardo-Lax *et al*. described a vector system combining an S-deleted SARS-CoV-2 replicon RNA complemented with an S-expressing plasmid (Ricardo-Lax et al., 2021). Both this team and Malicoat et al. (Malicoat et al., 2022) successfully pseudotyped their vectors with the G protein of the vesicular stomatitis virus (VSV-G) instead of S. Zhang *et al*. developed a similar system with ORF3 and E being the transcomplemented proteins, but their system did not include a marker gene (Zhang et al., 2021). Several studies showed that as expected, known pharmacological and biological inhibitors of SARS-CoV-2, such as remdesivir (RDV), type I interferons (IFN-I) and neutralizing antibodies, also efficiently reduced infection by SARS-CoV-2 vectors produced through complementation (Ju et al., 2021;Ricardo-Lax et al., 2021;Zhang et al., 2021;Malicoat et al., 2022). Thus, SARS-CoV-2 vector systems show great promise for use in antiviral screens. Yet, they still remain cumbersome to use, necessitating the repeated transfection of replicon RNA and of one or more plasmids for the expression of the complementing structural protein.

In this study, we successfully generated SARS-CoV-2 viral vectors in HEK293T cells constitutively producing SARS-CoV-2 VLPs made of all four structural proteins N, S, E and M through lentiviral transduction. To produce a vector able to transduce marker genes, these cells were electroporated with a SARS-CoV-2 replicon RNA deleted for S, M, and E and expressing a luciferase-GFP fusion protein. We show that expression of the marker genes following infection depends on replicon RNA replication and expression.

## Method

### Cell culture

SARS-CoV-2 N/S/E/M lentivirally transduced, virus-like particle (VLP)-producing HEK293T cells have been described before (Elfayres et al., 2023). To obtain human embryonic kidney HEK293T cells expressing ACE2 and TMPRSS2, HEK293T cells were stably transfected with plasmids expressing each of the human proteins along with a selectable marker. First, human TMPRSS2 cDNA was excised from the TMPRSS2 plasmid (Addgene #53887) (Edie et al., 2018) cut with BglII and Not1, and cloned into pcDNA3.1(+) Hygro cut with BamHI and NotI. HEK293T cells were co-transfected with linearized pcDNA 3.1(+) ACE-2 Flag (Genscript, Piscataway, NJ, USA) and pLVX puro (Takara Bio, Ann Arbor, MI, USA) using polyethylene imine (Syed et al., 2021) followed by selection with puromycin (1 ug/ml). One selected cell clone confirmed for ACE2 expression by Western blotting was transfected with linealized pcDNA 3.1(+) Hygro TMPRSS2 using PEI followed with hygromycin B selection (100 ug/ml). The HEK293T-ACE2/TMPRSS2 cell line was established from an isolated clone tested for TMPRSS2 expression. A549 adenocarcinomic human alveolar basal epithelial cells stably expressing human ACE2 and TMPRSS2 were generated as described before (Wang et al., 2023). Briefly, A549 cells (ATCC, CCL-185) were first transduced with lentiviral particles expressing human TMPRSS2. G418 (1 mg/ml) was added 40 h after transduction to select stably transduced cells. The selected cell population was further transduced with lentiviral particles expressing human ACE2, followed by selection with hygromycin B (1 mg/ml). Expression of ACE2 and TMPRSS2 was verified by Western blotting. Vero E6 African green monkey kidney epithelial cells (ATCC CRL-1586) were maintained in Eagle’s minimal essential medium (EMEM) supplemented with 10% fetal bovine serum and penicillin-streptomycin. All other cells were maintained in Dulbecco’s modified Eagle’s medium (DMEM) supplemented with 10% fetal bovine serum and penicillin-streptomycin. All cell culture media and supplements were from Cytiva Hyclone (Vancouver, Canada).

### Preparation of SARS-CoV-2 replicon-encoding bacterial artificial chromosome (BAC) DNA and *in vitro* replicon RNA transcription

The pSMART BAC v2.0 vector containing the SARS-CoV-2 (Wuhan-Hu-1) deleted in S, E and M and encoding a luciferase-GFP fusion (He et al., 2021) was obtained from the Biodefense and Emerging Infections Research Resources Repository (BEI resources; #NR-54972). To transform Stbl3 *E. coli* cells (Invitrogen, Burlington, ON, Canada) with the BAC construct, one vial of One Shot Stbl3 chemically competent cells was thawed on ice and mixed with 5 µL of DNA. The mixture was incubated on ice for 30 min followed by heat shock treatment at 42 °C for 45 sec without shaking. The vial was placed back on ice for 2 min, then 250 µL of pre-warmed SOC medium was added followed by incubation at 37°C for 1 h. The cells were plated onto LB plates containing 20 μg/mL chloramphenicol and cultured overnight at 37 °C. Individual colonies were isolated and grown, and then the BAC DNA was purified using the PhasePrep BAC DNA Kit (MilliporeSigma, Oakville, ON, Canada) according to the manufacturer’s instructions, and analyzed by restriction enzyme digestion.

For *in vitro* transcription, the BAC replicon-encoding DNA was first linearized with SwaI (New England Biolabs, Whitby, ON, Canada), and then purified by phenol/chloroform extraction followed by ethanol precipitation and resuspension in nuclease-free water. The mMESSAGE mMACHINE T7 transcription kit (Invitrogen) was used to generate replicon RNA from the linearized vector according to manufacturer’s instruction. Briefly, 180 μL of T7 transcription reaction, containing 7.2 μg of linearized BAC, T7 RNA polymerase as well as GTP, was incubated at 37 ^°^C for 2.5 h. 9 μL of TURBO DNase were then added and the reaction was incubated at 37 ^°^C for 15 min to degrade DNA. The resulting RNA was purified using the Monarch RNA cleanup kit (New England Biolabs) and analyzed by agarose gel electrophoresis.

### SARS-CoV-2 vector preparation

HEK293T cells continuously producing SARS-CoV-2 VLPs following stable transduction of S, N, E and M (Elfayres et al., 2023) were harvested using Trypsin/EDTA (Thermo Fisher Scientific, Mississauga, ON, Canada), washed with PBS, and resuspended in Neon Resuspension Buffer R (Invitrogen) to a final density of 1 × 10^7^ cells/mL. 30 μg of replicon RNA were added onto 2 × 10^6^ resuspended cells. The mixture (100 µL) was immediately transferred to Neon tips (Invitrogen), and electroporation was carried out in the Neon Device (Invitrogen) using the cell type-specific preloaded parameters (1350V, 20ms, 1 pulse). Transfected cells were transferred immediately into 10 cm plates containing prewarmed DMEM medium supplemented with 10% FBS and without antibiotics and placed in antibiotic-containing medium the next day. Viral vector-containing supernatants were collected 48 h post-transfection by centrifugation at 3,000 rpm for 10 min at 4^°^C. Supernatants were filtered through 0.45 μm PVDF filters (Millipore Sigma), loaded on top of 20% sucrose-PBS cushions and ultracentrifuged for 90 min at 30,000 rpm at 4^°^C. The pelleted particles were recovered in 50 μL of PBS solution and frozen at -80^°^C.

### Infections and flow cytometry

HEK293T-ACE2/TMPRSS2, A549-ACE2/TMPRSS2 and Vero E6 cells were plated in 96-well plates (3 × 10^4^ cells/well in 100 μL) in complete DMEM medium. After 24 h, cells were infected with 50 μL of concentrated SARS-CoV-2 viral vector preparation. 48 h later, cells were washed with 1X PBS and detached using 100 μL trypsin. The reaction was stopped by adding 150 µL of 4% formaldehyde diluted in PBS. The percentage of GFP-positive cells was determined by analyzing 20,000 cells on a CytoFLEX S flow cytometer (Beckman Coulter). Flow cytometry data were analyzed using FlowJo (Becton Dickinson). In some instances, remdesivir (Cayman Chemical, Ann Arbor, MI, USA) was added immediately after infection at a concentration of 100 nM. Virus titer (FACS infectious units/mL) was assessed by flow cytometry using the formula titer = ((F × Cn) /V) × DF, where F is the frequency of GFP-positive cells determined by flow cytometry, Cn is the total number of cells exposed to the vector preparation, V is the volume of inoculum (mL) and DF the dilution factor.

### Fluorescence microscopy

GFP expression in live cells was examined using an Axio Observer inverted fluorescence microscope (Zeiss) with a 20X objective, two days post-transfection with SARS-CoV-2 replicon RNA or two days post-infection with SARS-CoV-2 viral vectors. Images were recorded using the ZEN software.

### Luciferase assays

HEK293T-ACE2/TMPRSS2 cells were infected with the SARS-CoV-2 viral vector preparation as described above. Cells were processed for luciferase assay 48 h later. The Steady-Glo Luciferase Assay System kit from Promega (Madison, WI, USA) was used for all luciferase assays. Cells grown in 96-well plates were washed with PBS, trypsinized and plated in wells of a 96-well plate (black wall, clear bottom) at 100 µL per well. 100 µL of Steady-Glo Reagent were immediately added to each well. Luciferase activity was measured using the Biotek Synergy HT microplate reader according to the manufacturer’s instructions.

### SARS-CoV-2 vector infections in presence of interferons

Recombinant human IFN-α was obtained from Shenandoah Biotechnology (Warwick, PA). Recombinant human IFN-β and IFN-ω were obtained from PeproTech (Rocky Hill, NJ). HEK293T-ACE2/TMPRSS2 cells (3 × 10^4^ cells/well) were seeded in a 96-well plate in 100 μL complete DMEM medium. IFN-I was added to cell cultures 16 h prior to infection and at a final concentration of 20 ng/ml. Cells were next infected with 50 μL of SARS-CoV-2 viral vectors, as described above. Luciferase assays were performed 48 h post-infection as described above.

## Results

### Design and production of infectious SARS-CoV-2 viral vectors

The vector system described in this study was developed using HEK293T cells, which were modified by the simultaneous transduction of all four SARS-CoV-2 structural proteins (S, M, E, and N), as described in a previous report (Elfayres et al., 2023). As illustrated in Fig. 1A, we used both HEK293T cell populations continuously producing SARS-CoV-2 VLPs, as well as clones derived from these populations by limiting dilution (Elfayres et al., 2023). Cells were subjected to electroporation with an *in vitro*-produced SARS-CoV-2 replicon RNA (He et al., 2021). Following this, supernatants were harvested, and SARS-CoV-2 vector particles were purified and concentrated by filtration and ultracentrifugation. Subsequently, cells expressing ACE2 and TMPRSS2 were exposed to the vector preparations, and the resulting signals from GFP/luciferase activity were measured for quantification. As a preliminary test for SARS-CoV-2 vector production, HEK293T cells constitutively producing SARS-CoV-2 VLPs underwent electroporation with replicon RNA. Supernatants from the transfected cells were collected, concentrated and used to infect HEK293T-ACE2/TMPRSS2 cells. As illustrated Fig. 1B, GFP expression was detected by fluorescent microscopy both in the cells electroporated with the replicon RNA, as expected, and following infection of HEK293T-ACE2/TMPRSS2 with the supernatant. The GFP-positive transduced cells appeared healthy and dividing, suggesting that fluorescence was not an artifact from toxicity (Fig. 1B).

**Fig. 1.**
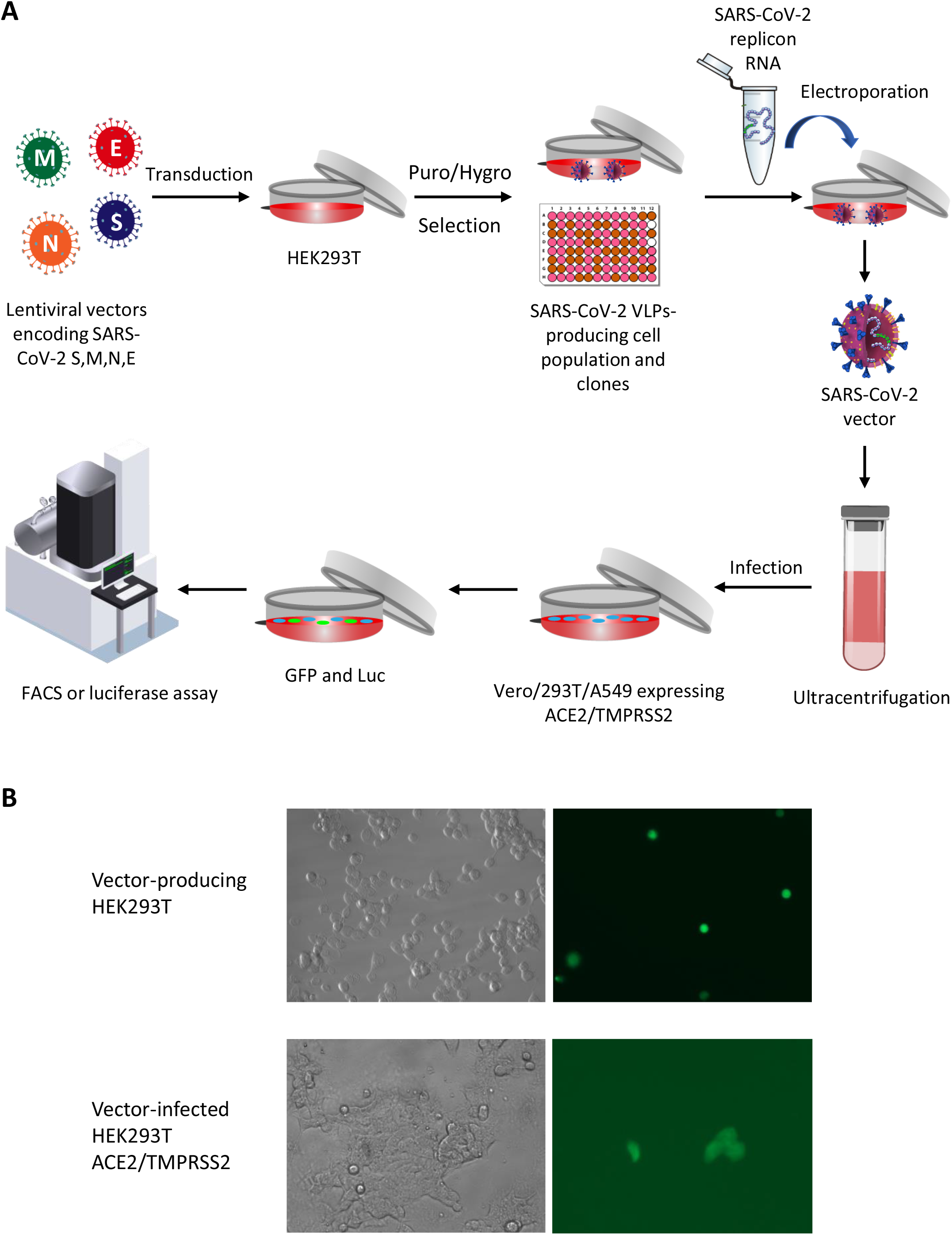
Design of a SARS-CoV-2 viral vector system through lentiviral transduction. (A) HEK-293T cells are transduced with lentiviral vectors encoding SARS-CoV-2 structural proteins to stably produce SARS-CoV-2 virus-like particles (VLPs). SARS-CoV-2 replicon RNA is subsequently electroporated into these cells. Vector particles are collected using low-speed centrifugation, followed by filtration and ultracentrifugation. The vectors are then used to infect SARS-CoV-2-permissive cell lines. Reporter gene expression (GFP and luciferase) is quantified using flow cytometry and luciferase assay, respectively. (B) Fluorescence microscopy analysis of HEK293T cells producing or infected with SARS-CoV-2 vectors. GFP detection in SARS-CoV-2 VLP-producing HEK293T cells following replicon RNA electroporation (top row), and in HEK293T cells expressing ACE2/TMPRSS2 following infection with SARS-CoV-2 vectors (bottom row).

We repeated this experiment, this time using a flow cytometer to quantify the proportion of infected cells (Fig. 2). In order to investigate the requirements for efficient vector production, we also tested the electroporation of the replicon RNA in cells co-transfected with all four SARS-CoV-2 structural proteins instead of being lentivirally transduced. We also electroporated the replicon RNA in two HEK293T cellular clones, 1-22 and 1-4, which were derived from the SARS-CoV-2 S/M/N/E transduced population (Elfayres et al., 2023). Finally, as an additional control, we also electroporated the SARS-CoV-2 replicon in parental HEK293T cells expressing no SARS-CoV-2 protein, or in cells expressing N only. GFP levels were measured by flow cytometry two days post-infection. The data show expression of GFP in 1.6% HEK293T-ACE2/TMPRSS2 cells upon infection with SARS-CoV-2 vectors prepared by electroporation of N/S/E/M-transduced cells with the replicon RNA (Fig. 2A). No GFP-positive cells were seen following infection with vector prepared by electroporation of the replicon RNA in untransduced cells or in cells transduced with N only, as expected. We did observe infection of HEK293T-ACE2/TMPRSS2 cells with SARS-CoV-2 vector produced in cells co-transfected with all four SARS-CoV-2 proteins along with the replicon RNA, but the frequency of transduced cells was lower (0.53%) compared to the vector produced from the N/S/E/M-transduced cells (1.61%). Thus, and at least based on these limited initial trials, the lentiviral transduction approach to express the SARS-CoV-2 structural proteins leads to higher vector yields. In addition, little to no GFP signal was detected in cells infected with SARS-CoV-2 vector in presence of remdesivir (0.01%), a nucleoside analog that abrogates replicon RNA replication (Beigel et al., 2020), showing that GFP expression was dependent upon replicon RNA replication and expression in the target cells. Based on the flow cytometry data obtained, and assuming no infectivity loss upon vector cleaning and concentration, we calculated a vector titer of 800 IU/mL for the unconcentrated supernatant from the N/S/E/M-transduced cells compared to 265 IU/ml from N/S/E/M-transfected cells. SARS-CoV-2 vectors produced in clone 1-22 and clone 1-4 yielded 0.84% and 0.27% GFP-expressing cells, respectively. The infectious titers for these two vector preparations were calculated at 420 IU/mL and 135 IU/mL, respectively. Results obtained with these two clones confirm that VLPs are being stably produced from the transduced cells, since establishment of those clonal populations required several weeks. However, the vector yields obtained were lower compared to the lentivirally transduced parental population.

**Fig. 2.**
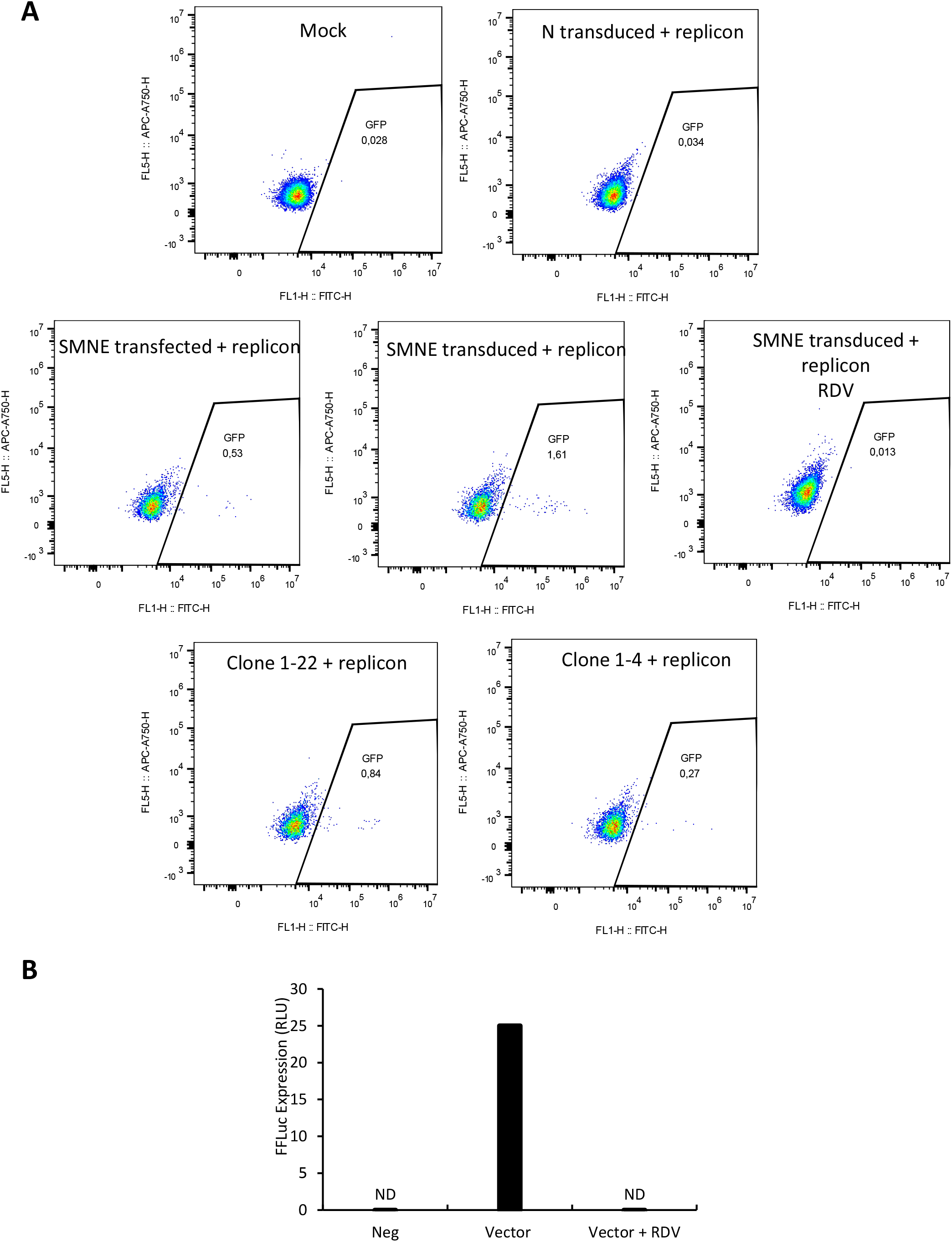
SARS-CoV-2 vectors transduce GFP and luciferase genes in ACE2/TMPRSS2-expressing HEK293T cells and exhibit remdesivir sensitivity. (A) HEK293T cells were exposed to SARS-CoV-2 vectors produced by electroporation of SARS-CoV-2 replicon RNA in a cell population stably transduced with S/M/N/E, or in selected clones (1-22, 1-4) derived from this population. As controls, (i) cells were exposed to the vector produced in S/M/N/E-transduced cells in the presence of 100 nM remdesivir; (ii) SARS-CoV-2 replicon RNA was electroporated in parental HEK293T cells (Mock) or in cells transduced with N only. GFP expression was quantified by flow cytometry 48 hours post-infection. (B) HEK293T-ACE2/TMPRSS2 cells were exposed to SARS-CoV-2 vectors and treated or not with remdesivir (RDV). Luciferase activity was measured 48 hours post-infection and is reported in relative light units (RLU), as described in the Materials and Methods section.

The SARS-CoV-2 replicon used in this study encodes a firefly luciferase-GFP fusion protein. Consequently, we anticipated a correlation between GFP expression and luciferase activity. HEK293T cells expressing ACE2 and TMPRSS2 were exposed to SARS-CoV-2 vector prepared in HEK293T cells lentivirally transduced with N/S/E/M, and luciferase activity was next quantified in cellular lysates. As shown in Fig. 2B, luciferase activity was detected post-infection, as evidenced by measurement of high RLU values compared to uninfected cells. Moreover, treatment with remdesivir resulted in a strong reduction in luciferase activity levels. Thus, these results recapitulate the GFP expression data, confirming the production of a viral vector expressing GFP and luciferase reporter proteins.

### SARS-CoV-2 viral vector challenges in different cell lines

To investigate the cellular tropism of the SARS-CoV-2 vector, A549 cells expressing ACE2/TMPRSS2 (Fig. 3A) and Vero E6 cells (Fig. 3B) were subjected to vector infection. Both cell lines were found to be permissive to infection (0.23% and 0.37%, respectively). The virus titer was 115 IU/mL and 185 IU/mL, as determined in these two cell lines. Fluorescence microscopy images showed healthy GFP-expressing cells, instead of signals attributable to autofluorescence induced by toxicity. Altogether, results from Fig. 2 and 4 suggest that this SARS-CoV-2 vector exhibits cellular tropism comparable to authentic SARS-CoV-2 virus.

**Fig. 3.**
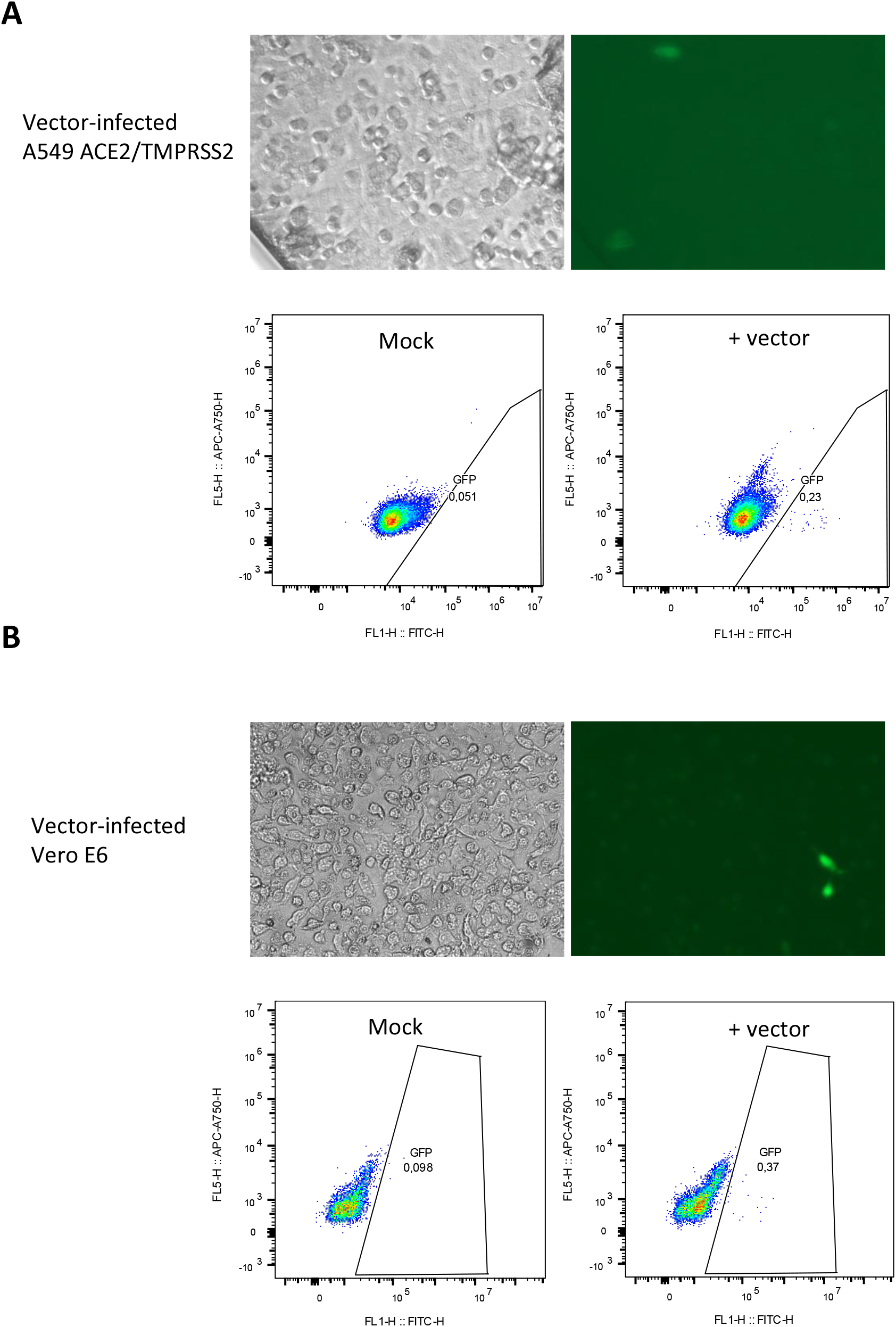
Detection of SARS-CoV-2 viral vector-mediated GFP expression in A549 ACE2/TMPRSS2 and Vero E6 cells. GFP detection in A549-ACE2/TMPRSS2 cells (A) or Vero E6 cells (B) infected with the SARS-CoV-2 vector, as analyzed by fluorescence microscopy and flow cytometry 48 hours after infection.

**Fig. 4.**
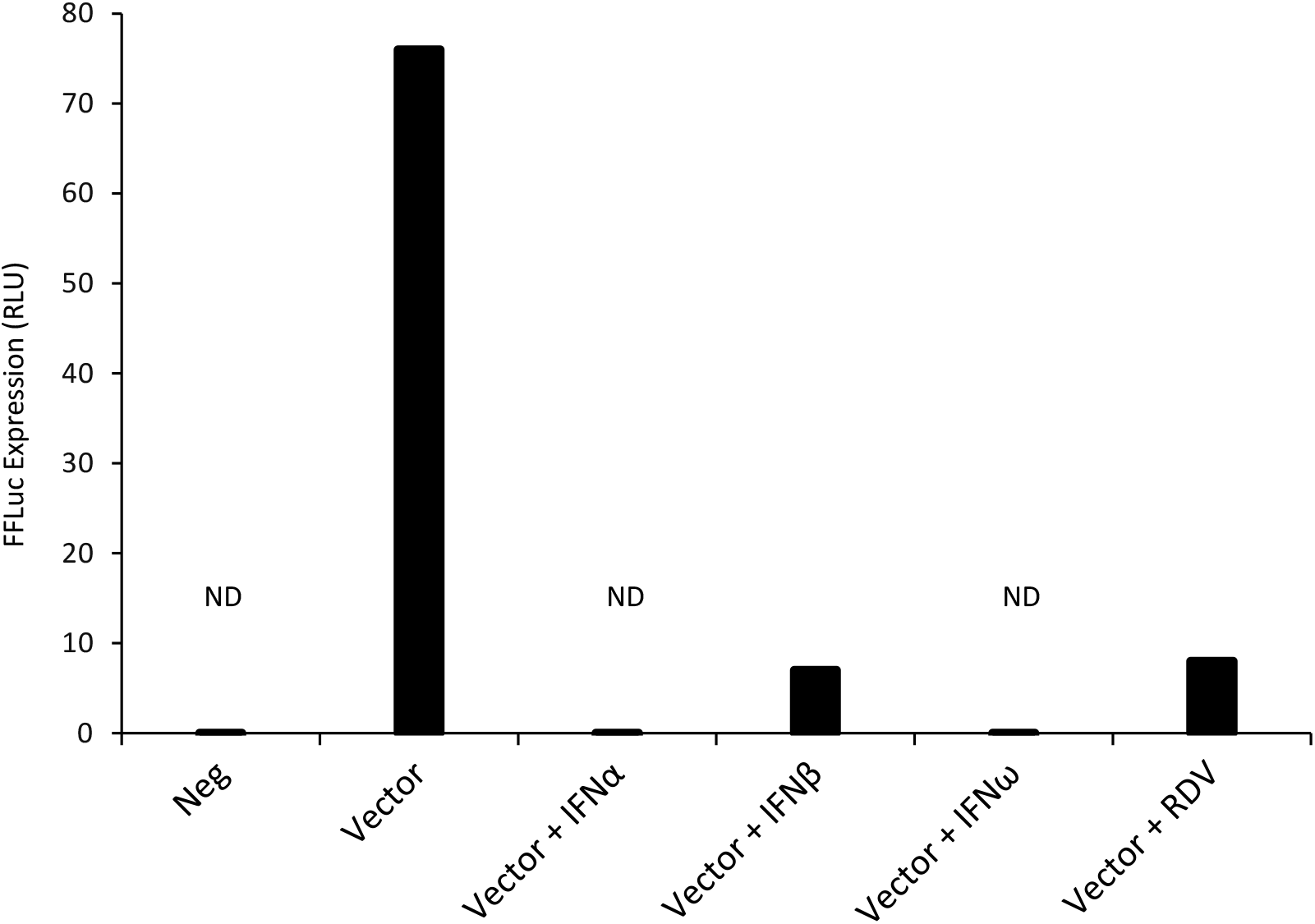
Inhibition of SARS-CoV-2 vector-mediated infection of HEK293T-ACE2/TMPRSS2 cells by type I interferons (IFN-I). HEK293T cells were pre-treated with IFN-α, IFN-β, or IFN-ω for 16 hours prior to infection with the SARS-CoV-2 vector. As a control, infection was also performed in the presence of 100 nM remdesivir. Luciferase activity was measured 48 hours post-infection.

### Inhibition of SARS-CoV-2-mediated infection by IFN-I

We employed the SARS-CoV-2 vector system to assess the antiviral efficacy of type I interferons (IFN-I), specifically human IFN-α and IFN-β, which are known for their capability to hinder SARS-CoV-2 infection, and IFN-ω, which to the best of our knowledge had not been tested yet. All three IFN-I subtypes efficiently decreased SARS-CoV-2 vector infectivity, as evidenced by luciferase assay (Fig. 4), without inducing visible cytotoxicity to target cells at the tested concentration. Moreover, we again noted SARS-CoV-2 vector inhibition by remdesivir. This result suggests that the SARS-CoV-2 vector described here has a potential use in SARS-CoV-2 antiviral drug screening.

## Discussion

Working with SARS-CoV-2 often requires strict containment measures in BSL-3 laboratories, which restricts research on the virus and the development of antiviral therapies. In this study, we developed a SARS-CoV-2 viral vector that offers a safer option for use in BSL-2 settings. The vector is based on SARS-CoV-2 replicon RNA lacking the structural protein genes S, M and E, allowing for amplification of a reporter gene without generating infectious SARS-CoV-2 particles. Instead, the viral structural proteins are continuously expressed as VLPs in transduced HEK293T cells (Fig. 1). Notably, the SARS-CoV-2 viral vector is capable of infecting two human cell lines that were modified to express the virus receptors ACE2 and TMPRSS2, replicating essential characteristics of the original virus, such as viral particle assembly, entry into ACE2-expressing cells, viral RNA replication and viral gene expression. To provide evidence that replicon RNA replication was required for efficient GFP/luciferase expression, remdesivir sensitivity was repeatedly tested and demonstrated to abrogate these signals. Additionally, in contrast to other SARS-CoV-2 vectors that have been previously reported, the one described here uses stable lentiviral transduction to express the four SARS-CoV-2 structural proteins, facilitating subsequent vector production. It should be noted, moreover, that this system probably constitutes a safer option compared to vectors in which the replicon has only one deleted structural protein (Ju et al., 2021). A key use of viral vectors is in screening potential drugs. The SARS-CoV-2 viral vector described here recapitulates all the different steps of the natural infection, making it a more accurate model for antiviral testing compared with previous systems (Syed et al., 2021). To illustrate this point, we show for the first time that SARS-CoV-2 infection is sensitive to IFN-ω (Fig. 4). Future research should focus on expanding the use of this vector system to high-throughput screening of novel therapeutic candidates targeting the structural proteins and replication machinery of SARS-CoV-2. However, a major limitation of this vector system as a drug screening tool is the low infectious titers. In this study, the detected vector titers ranged from 115 IU/mL to 800 IU/mL, which is significantly lower compared to the viral titers reported in other SARS-CoV-2 vector studies. One possible improvement could come from optimizing relative expression levels for the four structural proteins. For instance, vectors that use lower amounts of spike protein relative to other structural proteins were found to have higher titers (Liu and Liu, 2023). Another possible explanation is that the large size of the SARS-CoV-2 replicon RNA creates a bottleneck to replicon stability, electroporation and vector production. A potential solution to this problem could be to introduce mutations in N that enhance RNA expression, thereby promoting increased viral particle assembly (Syed et al., 2021). Another possible approach would be to test alternative SARS-CoV-2 replicons, particularly focusing on DNA-launched replicons (Szurgot et al., 2021;Malicoat et al., 2022;Liu and Liu, 2023). These replicons offer a promising route for enhancing efficiency and consistency in viral production. By integrating the replicon directly into the host cellular genome, cells could be engineered to stably express the replicon continuously, bypassing the need for repeated RNA electroporations. This would not only improve efficiency but also reduce variability, as stable expression systems tend to generate more consistent viral titers over time (Su et al., 2023). Establishing stable cell lines would also open up opportunities for scalable production, essential for drug screening, vaccine development, or other therapeutic applications where high and consistent viral titers are critical. Higher vector titers will also be necessary to explore infections in animal models. A promising approach would involve testing this vector in SARS-CoV-2-permissive mouse models, to study viral tropism and the capacity of the vector to infect airway cells following delivery in aerosolized form (Knight et al., 2021). This, in turn, would open the door to using a SARS-CoV-2-derived vector in vaccine development.

## Conclusions

This study contributes advances toward the ultimate goal of developing DNA-based, stable, scalable coronaviral vector production systems, with applications in anticoronaviral drug discovery as well as vaccine development.

## Author contributions

L.B. and G.E. conceptualized the study; G.E. performed the analyses; G.E. and L.B. interpreted the data; Y.X., Q.P., C.L. and B.B. provided essential materials; G.E., L.B. and B.B. drafted the manuscript; all authors have read and approved the final version of the manuscript.

